# The termite fungal cultivar *Termitomyces* combines diverse enzymes and oxidative reactions for plant biomass conversion

**DOI:** 10.1101/2021.01.13.426627

**Authors:** Felix Schalk, Cene Gostinčar, Nina B. Kreuzenbeck, Benjamin H. Conlon, Elisabeth Sommerwerk, Patrick Rabe, Immo Burkhardt, Thomas Krüger, Olaf Kniemeyer, Axel A. Brakhage, Nina Gunde-Cimerman, Z. Wilhelm de Beer, Jeroen S. Dickschat, Michael Poulsen, Christine Beemelmanns

## Abstract

Macrotermitine termites have domesticated fungi in the genus *Termitomyces* as their primary food source using pre-digested plant biomass. To access the full nutritional value of lignin-enriched plant biomass, the termite-fungus symbiosis requires the depolymerization of this complex phenolic polymer. While most previous work suggests that lignocellulose degradation is accomplished predominantly by the fungal cultivar, our current understanding of the underlying biomolecular mechanisms remains rudimentary. Here, we provide conclusive OMICs and activity-based evidence that *Termitomyces* partially depolymerizes lignocellulose through the combined actions of high-redox potential oxidizing enzymes (laccases, aryl-alcohol oxidases and a manganese peroxidase), the production of extracellular H_2_O_2_ and Fenton-based oxidative degradation, which is catalyzed by a newly described 2-methoxybenzoquinone/hydroquinone redox shuttle system and mediated by secreted chelating dicarboxylic acids. In combination, our approaches reveal a comprehensive depiction of how the efficient biomass degradation mechanism in this ancient insect agricultural symbiosis is accomplished through a combination of white- and brown-rot mechanisms.

**Importance:** Fungus-growing termites have perfected the decomposition of recalcitrant plant biomass to access valuable nutrients by engaging in a tripartite symbiosis with complementary contributions from a fungal mutualist and a co-diversified gut microbiome. This complex symbiotic interplay makes them one of the most successful and important decomposers for carbon cycling in Old World ecosystems. To date, most research has focused on the enzymatic contributions of microbial partners to carbohydrate decomposition. Here we provide genomic, transcriptomic and enzymatic evidence that *Termitomyces* also employs redox mechanisms, including diverse ligninolytic enzymes and a Fenton-based hydroquinone-catalyzed lignin-degradation mechanism, to break down lignin-rich plant material. Insights into these efficient decomposition mechanisms open new sources of efficient ligninolytic agents applicable for energy generation from renewable sources.

## Introduction

Among the different types of nutritional symbiosis, crop agriculture represents one of the most sophisticated systems. Beyond examples from humans, only a few insect lineages maintain and manure external symbiotic partners.^1^ Fungus-growing termites (*Macrotermitinae*) underwent a major transition ca. 30 Mya when they started to domesticate the mutualistic fungus *Termitomyces* (Agaricales, Lyophyllaceae) as their main food source.^2,3^ Since then, fungus-growing termites have become major biomass decomposers of dead plant material resulting in a substantial ecological footprint in the Old World (sub)tropics.^4,5^

*Termitomyces* is manured by termite workers in a cork-like structure termed the “fungus comb”, which is found within in the underground chambers of the termite mound and is comprised of predigested plant material (Figure 1A, B).^6^ Old termite workers collect and transport the necessary plant material while younger workers macerate and ingest the plant material along with asexual *Termitomyces* spores and enzymes, which are produced in fungal nodules on the mature parts of the fungal comb.^1,2,7^ The resulting lignocellulose and spore-enriched feces are then used to craft fresh fungus comb. After spore germination, the fungus matures within 15-20 days and energy-rich fungal nodules are formed to serve as the major food source for younger workers.^8^ After an average turn-over time of 45-50 days the remains of the comb material serve as the major nutrition of older workers resulting overall in the nearly waste-less decomposition and recycling of plant material.^9^

**Figure 1.**
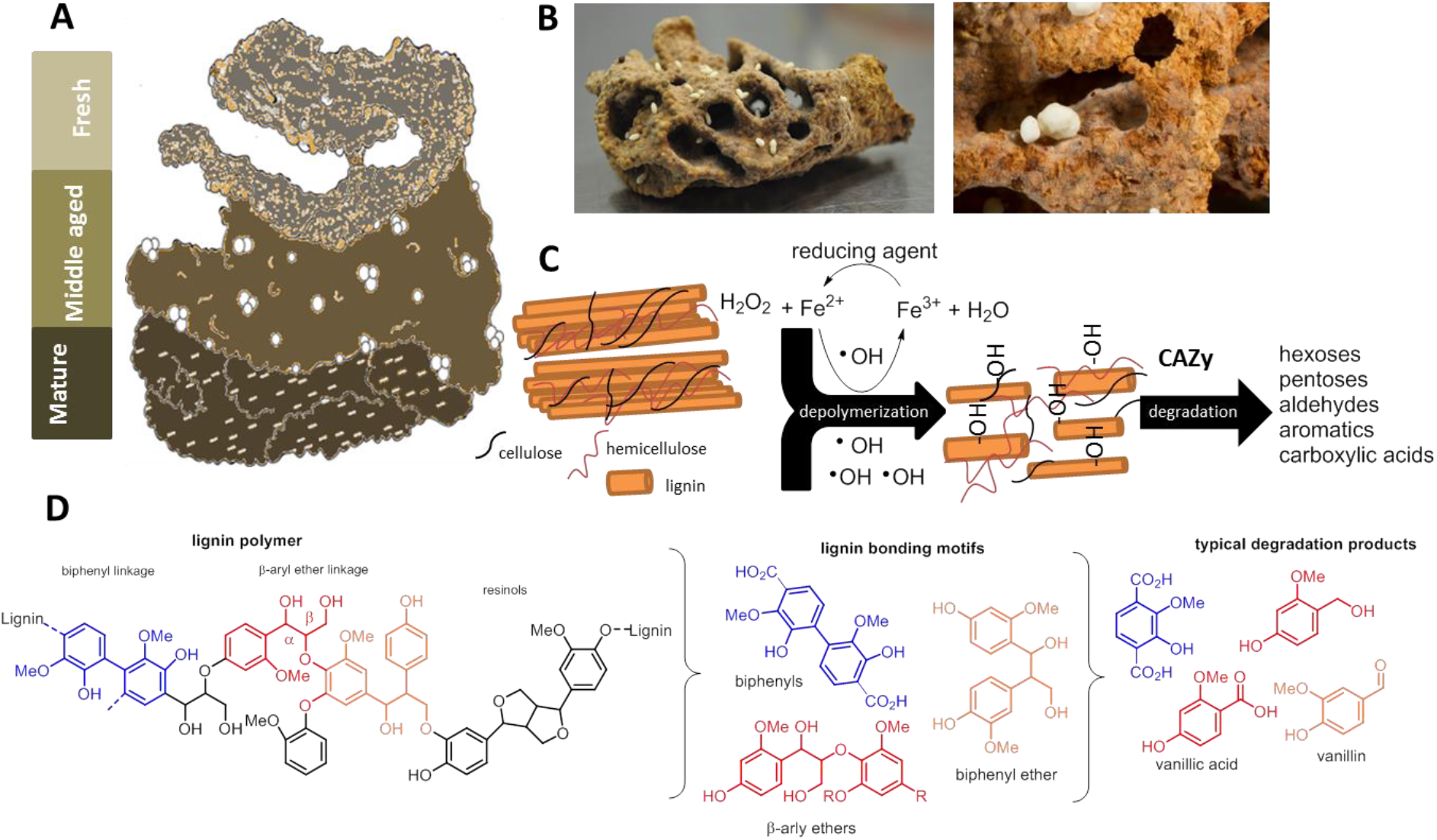
A) Schematic representation of fungus comb at different maturation stages. B) Freshly collected mature fungus comb carrying fungal nodules. C) Schematic representation of lignin depolymerization via hydroxylation and oxidative cleavage with subsequent degradation by CAZy enzymes to smaller metabolites. D) Schematic structure of lignin biopolymer, lignin motifs and lignin-derived degradation products.

Although the feeding behavior of termites has been studied in detail for decades,^10^ the biochemical mechanisms for degrading the foraged plant biomass has remained largely unresolved and a topic of intensive discussion.^1,11^ Plant biomass consists mostly of lignocellulose, a complex matrix consisting of cell wall polysaccharides: cellulose (40-50%), hemicellulose (25-30%), and the structurally complex and inhomogeneous phenolic polymer lignin (15-20%).^12^ The depolymerization and degradation of lignin provides an enormous energetic burden to any microorganism due to its inhomogeneous nature, and the strong covalent carbon-carbon and carbon-oxygen linkages between hydroxycinnamoyl alcohol derived monomers that are covalently cross-linked to plant polysaccharides (Figure 1C, D).^13,14^ However, once oxidative mechanisms have broken up the dense lignin structure, degrading enzymes are able to diffuse into the material and access valuable embedded biphenylic, phenolic and carbohydrate reservoirs.^15,16,17^ Although the degradation process appears to be a necessary endeavor to manure the complex fungus-termite-bacteria symbiosis, the fate of lignin within termite fungus combs still remains unclear.

A recent study on fungus comb pretreatment in *Odontotermes formosanus* by Li *et al.* indicated that lignin is partly cleaved during the first gut passage.^18^ Additionally, it was hypothesized that *Termitomyces* might have lost key delignification potential throughout its evolutionary history with the termites. However, previous and more recent transcriptomic and analytically-guided studies in other Macrotermitinae species by Poulsen and coworkers showed that fresh comb from *Odontotermes* spp. and *Macrotermes natalensis* is lignin rich,^7^ suggesting that the role of gut passage in lignin cleavage may differ between termite species.^9^ Based on fungal RNAseq analysis and enzymatic assays, the study reasoned that maturation of the fungus comb causes the decomposition of the lignocellulose-rich biomass through the actions of fungal and/or bacterial enzymes.

These partially contradictory results led us to investigate whether *Termitomyces* has the capacity to depolymerize or even degrade lignin-rich biomass. Hence, we commenced our analysis by comparative genome analysis of nine *Termitomyces* species and assessment of their capacity to produce ligninolytic enzymes (e.g., laccase (Lac), lignin peroxidase (LP), manganese peroxidase (MnP), and/or versatile peroxidase (VP)) and enzymes supporting degradative pathways (e.g., aryl-alcohol oxidase and quinone reductases).^19,20^ Here, we show that *Termitomyces* has the capacity to produce a broad diversity of laccases and a MnP similar to other basidiomycetes, but lacks other necessary class II peroxidases (e.g., LPs and VPs) required for the complete degradation of non-phenolic lignin as is known from other basidiomycete white-rot fungi.^13,14,16^ These findings were supported by analysis of gene expression levels in RNAseq datasets of fungus combs at different maturation stages. Additional *in silico* and biochemical studies led us to the conjecture that *Termitomyces* might employ hydroquinone-mediated Fenton chemistry (Fe^2+^ + H_2_O_2_ + H^+^ → Fe^3+^ + ^•^OH + H_2_O) using a herein newly described 2-methoxy-1,4-dihydroxybenzene (2-MH_2_Q, **19**) based electron shuttle system to complement enzymatic lignin degradation pathways. We further deduced that the presence of small dicarboxylic acids produced by *Termitomyces* not only allows the fungus to solubilize necessary metal ions, but also mediates Fenton-based redox chemistry, making the system one of the most successful farming insect symbioses.

## Results

### Genomic and transcriptomic analysis of lignocellulolytic capacity

First, we subjected two *Termitomyces* species, excavated in South Africa in 2011 and 2015, to whole genome sequencing using Illumina sequencing technology (LGC Genomics (Berlin, Germany)) and RNA sequencing using the BGISeq-500 platformg (BGI, Hong Kong). Annotated genomes of both species were obtained using Augustus 3.3.3 after RNAseq data was mapped to the genomes and used for algorithm training. The resulting draft genome of *Termitomyces* sp. T153 (*Macrotermes natalensis*) had an estimated size of 84.1 Mb (scaffold N50 = 23.88 kb) with more than 13,000 genes (Accs. Nr. JACKQL000000000). Similarly, the draft genome of *Termitomyces* sp. T112 (*Macrotermes natalensis*) had an estimated size of 79.8 Mb (scaffold N50 = 33.34 kb) and also >13,000 genes (Accs. Nr. JACKQM000000000). For further analysis, we also re-annotated seven *Termitomyces* genomes deposited at GenBank, including our previously reported *Termitomyces* sp. J123 (alias P5) from *Macrotermes natalensis*,^3^ using the same settings in Augustus 3.3.3 (for details, see doi:10.5281/zenodo.4431413: Table S2 and S3). To gain insights into the functional capacity for biomass degradation, we first identified CAZyme families within each genome using a local installation of the dbCAN2 server.^21,22,23^ As depicted in Figure 2, comparison of all nine *Termitomyces* genomes revealed comparable numbers of polysaccharide-degrading enzymes, such as exo-cellobiohydrolases, endoglucanases assigned to different glycoside hydrolase families (GH) and lytic polysaccharide monooxygenase (LPMOs), with no particular enrichment or reduction of CAZy families compared to basidiomycete reference genomes (for details, see doi:10.5281/zenodo.4431413: Figure S1-S3).^24^ We also searched *Termitomyces* genomes for the presence/absence of gene sequences encoding for highly oxidizing proteins that could contribute to the depolymerization and catabolic degradation of lignin,^18^ and contained on average 16 gene sequences encoding for laccases (AA1, EC 1.10.3.2),^25,26,27,28^ oxidases that use diphenols and related substances as electron donors and oxygen as the acceptor, thereby creating reactive C- and O-based radical species in the process. In addition, we identified a putative manganese peroxidase (MnP, AA2, EC 1.11.1.13), which generates redox-active Mn^3+^ species, and a subset of alcohol oxidases and dehydrogenases (AA3 and AA5) that catalyze the oxidation of (aryl) alcohols or carbohydrates with the concomitant formation of hydroquinones and/or H_2_O_2_ that could be used by other peroxidases.^29,30^ We also identified iron reductase domains (AA8) and putative benzoquinone reductases (AA6) that are key to maintain efficient Fenton-chemistry-based redox cycles by reductive Fe^2+^ sequestration and regeneration of organic benzoquinone-based redox shuttles. However, all investigated *Termitomyces* genomes lacked signs of the class II peroxidases (e.g., LPs and VPs) that are normally found in white-rot fungi and necessary for the enzymatic mineralization of lignin (for details, see doi:10.5281/zenodo.4431413: Table S4-S12).^31^

**Figure 2.**
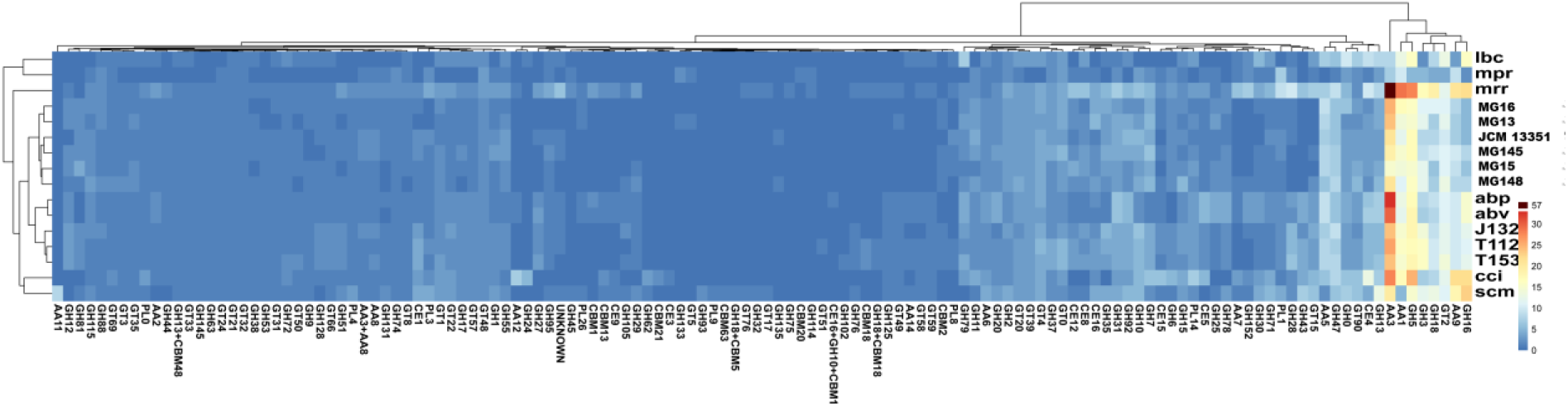
A) Heatmap of the numbers of hits for representatives of different CAZy families in the predicted proteomes of *Termitomyces* spp. (T112, T153, J132, JCM 13351, MG145, MG16, MG15, MG148, MG13) and other selected basidiomycete fungi (*Laccaria bicolor* (lbc), *Moniliophthora perniciosa* (mpr), *Moniliophthora roreri* (mrr), *Agaricus bisporus var. Burnettii* (abp), *Agaricus bisporus var. Bisporus* (abv), *Coprinopsis cinerea* (cci), *Schizophyllum commune* (scm)). Vertical axis shows clustering of enzymes based on expression levels.

We subsequently analyzed the expression levels of candidate genes related to lignin depolymerization in RNAseq data obtained from three regions in the fungus comb (Figure 3):^7^ fresh comb, within which most plant-biomass decomposition is likely to occur; old comb where decomposition might still occur but to a lesser extent; and nodules, which feed young workers and serve as fungal spore and enzyme reservoirs (for details, see doi:10.5281/zenodo.4431413: Table S25). As depicted in Figure 3, we found transcription levels of genes encoding oxidative enzymes (e.g., Lac, MnP, AA3 and AA5) and enzymes that protect against reactive intermediates (e.g., benzoquinone reductase, super oxide dismutase, glutathione peroxidase, and peroxiredoxin) across all three datasets.

**Figure 3.**
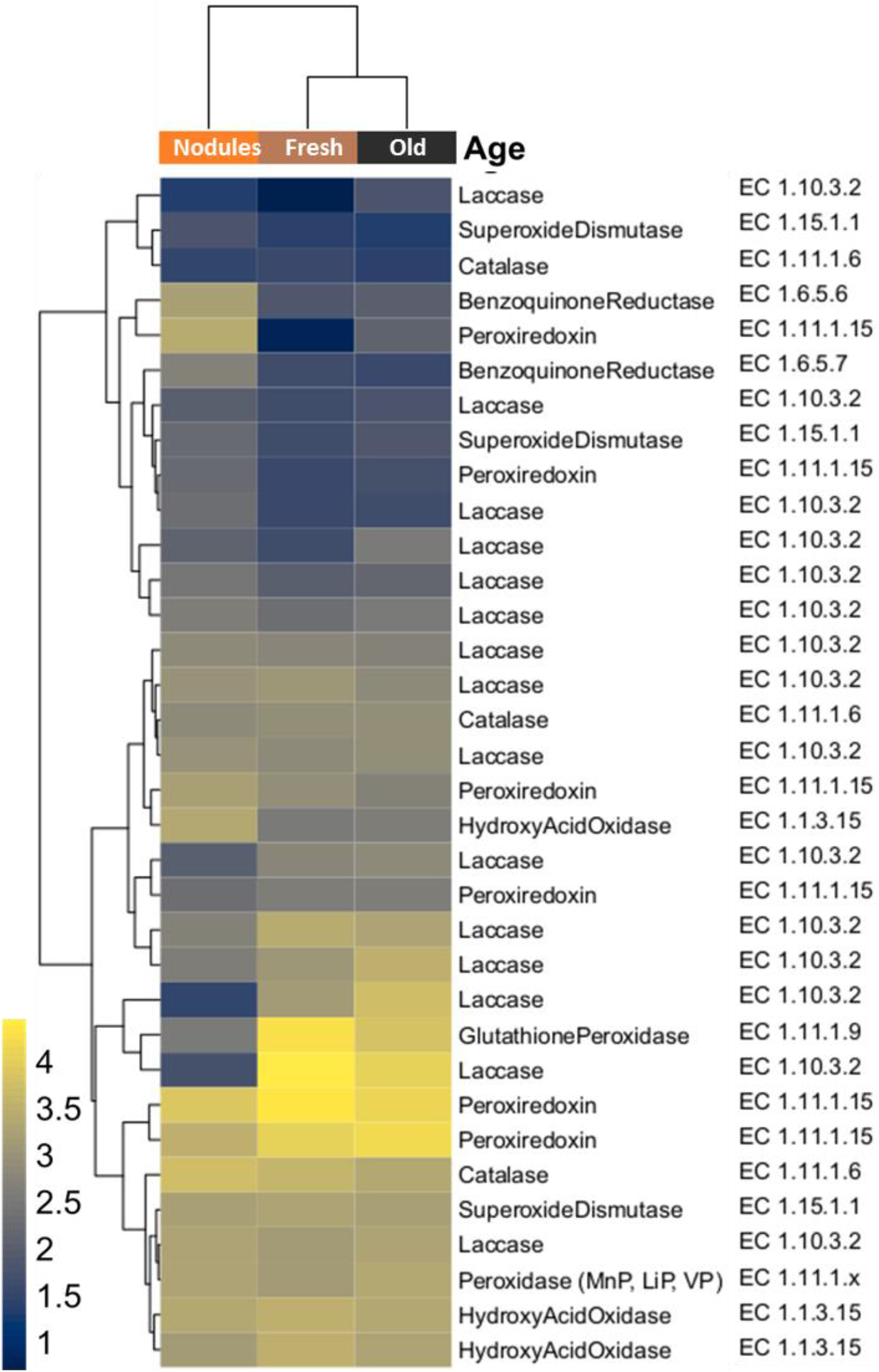
Heatmap of redox enzyme transcription levels based on RNAseq data of fresh comb, old comb and fungal nodules from *Macrotermes* colony Mn156.^7^ Transcript abundances are depicted as *log10 gene expression values* and color schemes were generated by “viridis” (Version 0.5.1).^32,33^

This genetic and transcriptomic survey revealed that *Termitomyces* has the genomic capacity to produce lignocellulolytic enzymes similar to other basidiomycetous fungi and may even be able to induce and catalyze Fenton chemistry,^34^ but lacks LiP, VP and other generic peroxidases that are needed to degrade the more recalcitrant non-phenolic components of lignin.^7^

### Fenton Chemistry of *Termitomyces*

Fenton chemistry involves the reaction between Fe^2+^ and H_2_O_2_ yielding Fe^3+^ and highly reactive hydroxyl radical (^•^OH), a powerful oxidant (*E^0^* = 2.8 versus normal hydrogen electrode) that is able to unselectively oxidize hydrocarbons and non-phenolic aromatic units within lignocellulose-rich material. Brown-rot fungi are known to make use of Fenton chemistry to depolymerize lignocellulose biomass^35^ and modulate the redox potential of Fe^2+/3+^ species by secretion of dicarboxylic acids that act as chelators to form diffusible Fe-complexes and as proton donors for catalytic degradation processes.^36^ Additionally, redox-active fungal quinones (Q) and hydroxyquinones (H_2_Q), such as 2,5-dimethoxy-1,4-benzoquinone (2,5-DMQ), 2,5-dimethoxy-1,4-hydroquinone (2,5-DMH_2_Q), and its regioisomer 4,5-dimethoxy-1,2-benzendiol (4,5-DMH_2_Q)), have been discussed to serve as redox shuttles (3H_2_Q + 2O_2_ → 3Q + 2H_2_O + 2HO^•^) in Fenton-chemistry of rotting fungi (e.g. *S. lacrymans*, the *Gloeophyllales* and the *Polyporales*)^37,38,39^ as they have the ability to switch between oxidation states via one-electron transfer reactions that allows for the concomitant formation of Fe^2+^ from Fe^3+^ and hydroxyl radicals (HO^•^) from O_2_ (Figure 5, 6).

Thus, we evaluated if *Termitomyces* employs any of those measures to enable lignin depolymerization by using *Termitomyces* sp. T153 and P5 as model strains. First, we employed a standardized colourimetric ferrozine assay to determine if extracellular Fe^3+^ is reduced to Fe^+2^ within the surrounding mycelium; a prerequisite to initiate Fenton chemistry.^40,41^ As depicted in Figure 4A, topical application of a ferrozine solution caused a clear color change within minutes, which was indicative for the immediate reduction of Fe^3+^ to Fe^+2^. Next, we determined the pH value within the fungal mycelium as enzyme activities, redox potential of H_2_O_2_ and metal complexes are strongly pH-dependent.^34^ Here, we found that *Termitomyces* acidifies the surrounding medium to as low as pH 5 (Figure 4D), which lies within the range of optimal enzyme activities of many lignin-degrading enzymes (pH 4.5-5.0).^14,20^ As the Fenton reaction also requires H_2_O_2_, we tested if *Termitomyces* generates sufficient extracellular H_2_O_2_ to initiate the reaction. Based on a H_2_O_2_-dependent colorimetric assay we found that *Termitomyces* generates approximately 4-6 μg extracellular H_2_O_2_ per gram fungal mycelium during growth on solid support (mycelium age: 7-21 days, for details, see doi:10.5281/zenodo.4431413: Table S17, S18).

**Figure 4.**
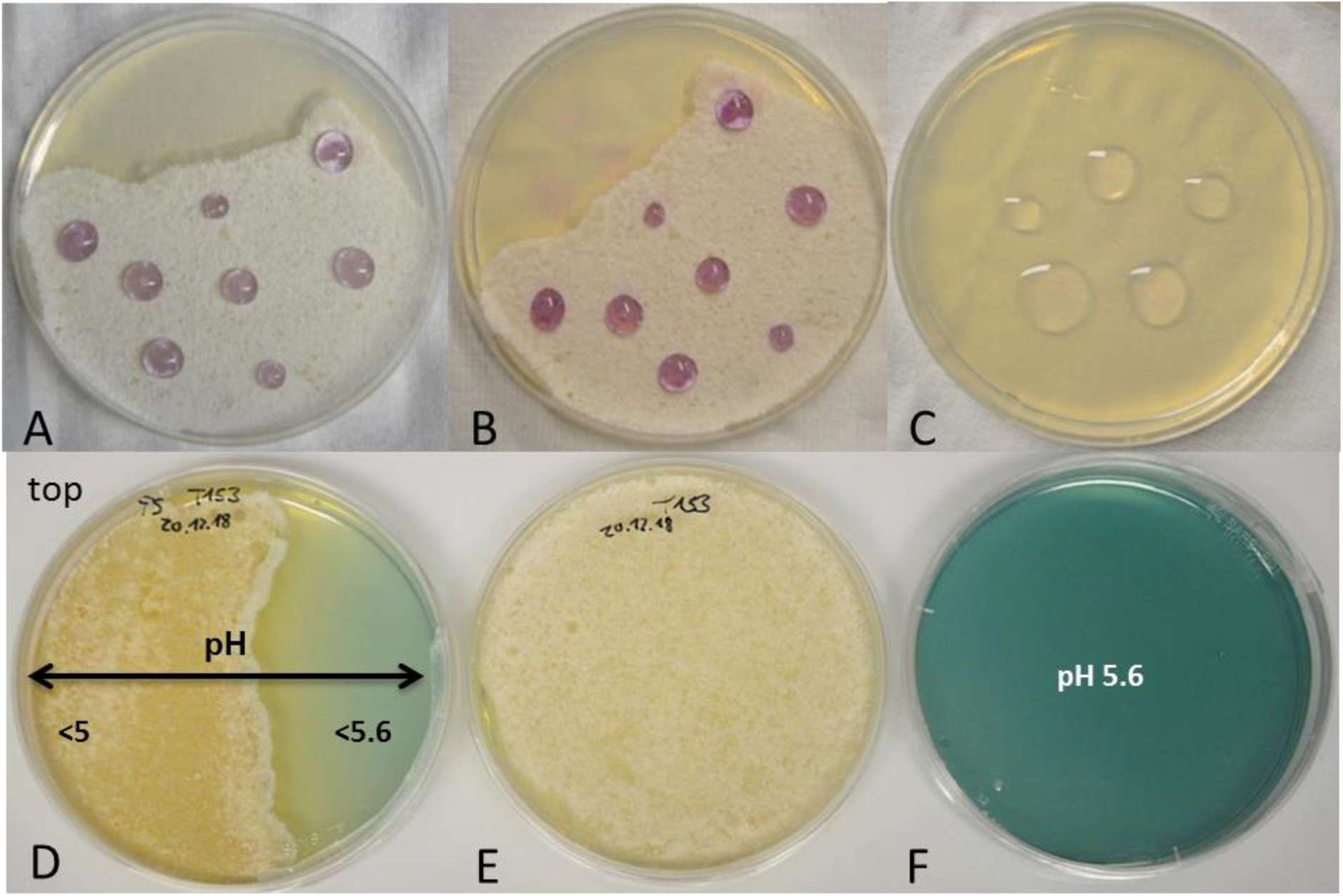
A) Ferrozine solution added to a *Termitomyces* sp. T153 culture grown on PDA (18 d) and incubated for 5 min and B) 30 min; C) ferrozine solution on PDA plate (negative control); D) *Termitomyces* sp. T153 grown on PDA containing D) bromocresol green as pH indicator (day 28) and E) without indicator; F) PDA plate containing bromocresol green.

**Figure 5.**
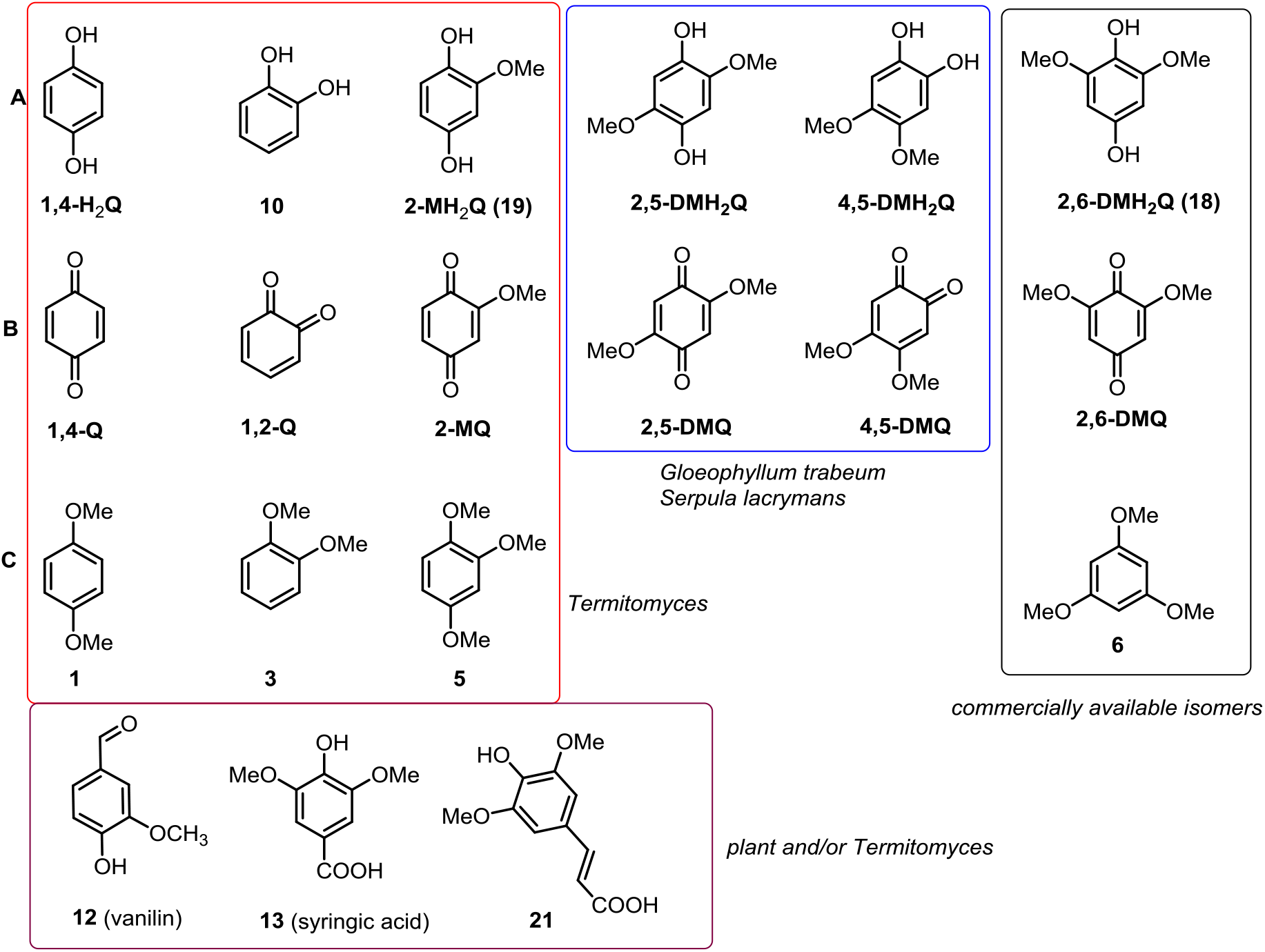
Structures of redox active compounds discussed in this work. A) Hydroxyquinones (H_2_Q), B) corresponding quinones (Q) and C) methoxylated derivatives of H_2_Q. Compounds identified from *Termitomyces* are highlighted in a redox box, compounds identified from other rotting fungi are marked with a blue box, derivatives isolated from *Termitomyces* and of plant origin are highlighted in a purple box and commercial derivatives for comparison are highlighted in a black box.

**Figure 6.**
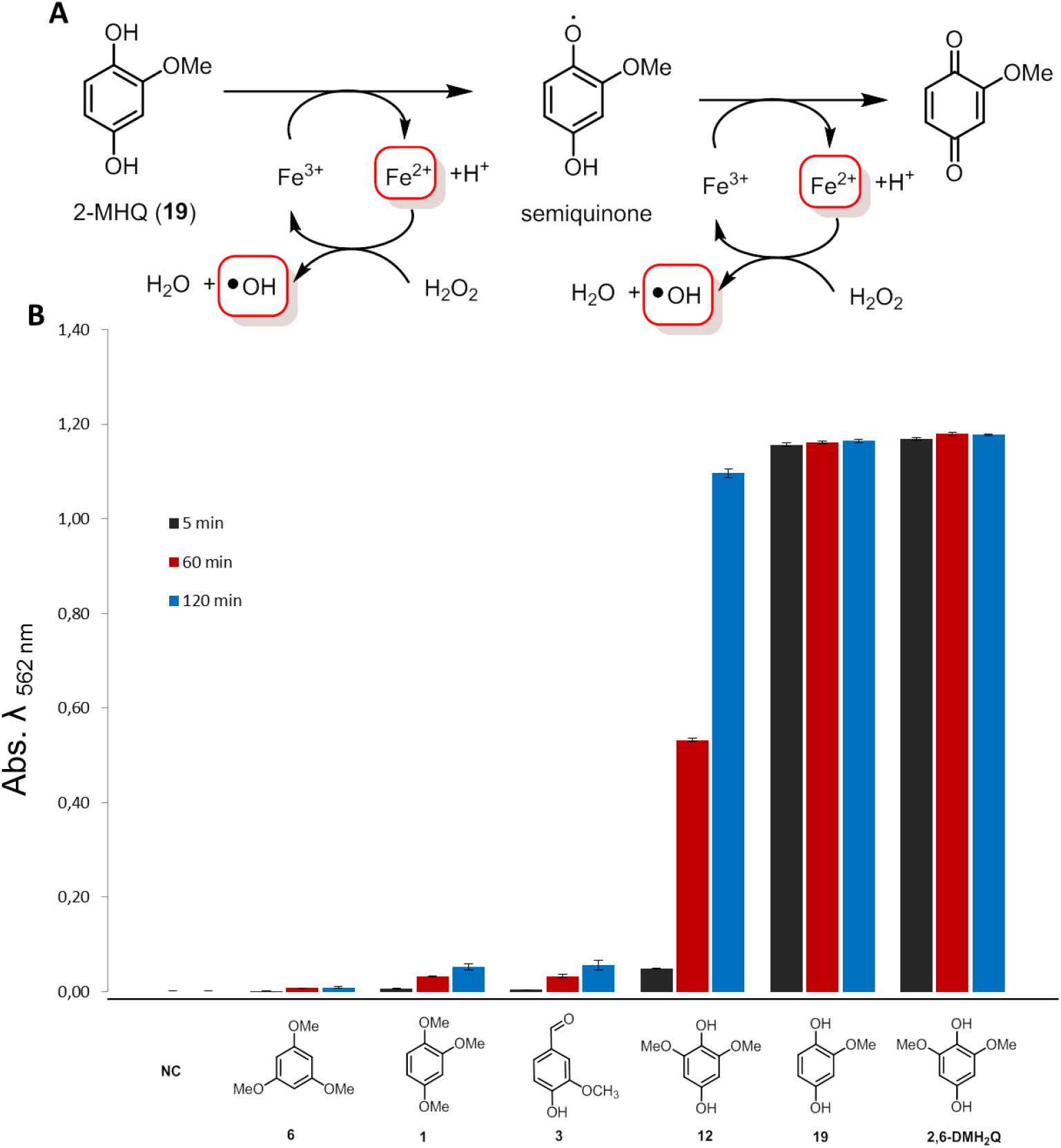
A) Mechanistic depiction of the 2-MH_2_Q initiated Fenton-reaction via the formation of a radical semiquinone species and oxidation to 2-MQ; B) quantification of Fe^3+^ reduction by H_2_Q using a colorimetric Ferrozine-based assay (NH_4_OAc buffer, pH = 4).

In a next step, we evaluated if *Termitomyces* produces redox-active redox-active H_2_Q/Q using gas chromatography coupled with mass spectrometry (GC-MS). Although the formation of previously reported 2,5-DM(H_2_)Q was not observed, we were intrigued to detect 2-methoxy-1,4-benzoquinone (2-MQ), its reduced H_2_Q named 2-methoxy-1,4-dihydroxybenzene (2-MH_2_Q) and the fully methylated derivative 1,2,4-trimethoxybenzene (**5**), as well as other structurally related (di)methoxylated hydroxybenzenes (e.g. **1**, **3**, **12**) (Figure 5). We also verified the identity of the newly detected quinone derivatives 2-MQ and 2-MH_2_Q by synthesis and comparison of GC-MS retention times (for experimental details, see doi:10.5281/zenodo.4431413: Figure S21, S22, Table S14, S15, S22).

We then evaluated the ability of H_2_Qs to reduce Fe^3+^ to Fe^2+^ using the established Ferrozine-based Fe^3+^-reduction assay.^42^ Overall, 2,6-DMH_2_Q (**18**), a regioisomer of 2,5-DMH_2_Q was the most reactive derivative that was able to reduce Fe^3+^ to Fe^2+^ within seconds, and was therefore used as a positive control in further experiments (Figure 6). In comparison, 2-MH_2_Q (**19**) showed a slightly reduced activity, which is likely a reflection of the electronic effect caused by the lack of one additional electron-donating -OCH_3_ group. We also tested the reducing ability of other (methoxylated) hydroxybenzenes, all of which showed a reduced reactivity compared **18** and **19**. Subsequently, we expanded our studies to combinations of redox active derivatives and were able to observe in most cases the superposition of redox activities (for experimental details, see doi:10.5281/zenodo.4431413: Figure S9, Table S19)

As Fenton chemistry produces highly reactive hydroxyl radical (•OH) we then confirmed the presence of these short-lived radicals in our H_2_Q-mediated Fenton reactions using a fluorometric assay based on the reaction with terephthalic acid (TPA). Similar to literature reports for 2,6-DMH_2_Q (**18**),^36,39^ the newly identified and structurally related H_2_Q **19** catalyzed the formation of ^•^OH in the presence of H_2_O_2_ and Fe^3+^ within seconds. In contrast, derivatives such as 1,2-dihydroxybenzene (**10**) and syringic acid (**13**) caused formation of hydroxyl radicals with lower initial reactivity but over a period of more than 90 min (for details, see doi:10.5281/zenodo.4431413: Figure S6). Having verified that *Termitomyces* produces reactive H_2_Qs that are able to induce the formation of Fenton reagents (Fe^2+^, H_2_O_2_ and ^•^OH), we then elaborated on the influence of fungal-derived dicarboxylic acids (oxalic acid, tartaric acid, malic acid, fumaric acid and succinic acid) ^43,44^ on the Fenton reaction (Figure 7). While low concentrations of oxalic acid (0.1 mM) influenced the reducing ability of H_2_Qs only mildly, increasing concentrations started to abolish their reducing capability in a concentration dependent manner with only the most reactive 2,6-DMH_2_Q (**18**) able to reduce Fe-oxalate complexes in the presence of less than 5.0 mM oxalic acid (for details, see doi:10.5281/zenodo.4431413: Figure S11-S13).^45^ At 10 mM oxalic acid a significant amount of autoxidation-related Fe^3+^-reduction was observed. A similar trend was observed for tartaric acid as a chelating agent, albeit with a stronger autoxidation effect.^46^ In contrast, malic, fumaric and succinic acid only moderately altered the redox potential and showed minor tendencies towards autoxidation. The overall ability of H_2_Q to reduce dicarboxylic acid complexes of Fe^3+^ decreased in the following order: oxalic acid > tartaric acid > malic acid >> fumaric acid ≥ succinic acid (for details, see doi:10.5281/zenodo.4431413: Figure S14-S16).

**Figure 7.**
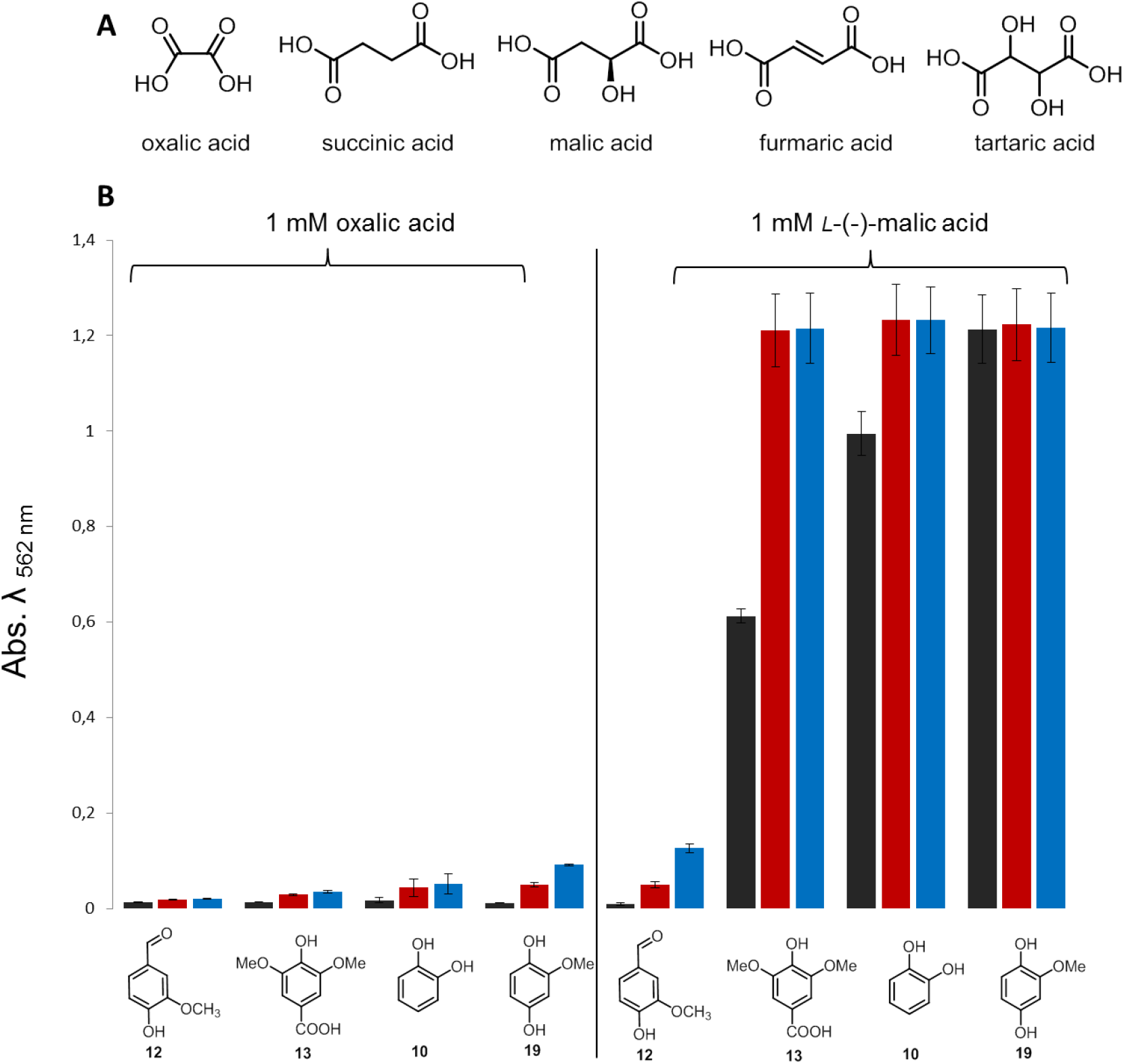
A) Structures of metal-chelating dicarboxylic acids; B) quantification of Fe^3+^ reduction by H_2_Q using a colorimetric Ferrozine-based assay in in the presence of 1 mM oxalic acid and 1 mM malic acid.

While laboratory culture conditions generally supply sufficient Fe-concentrations for growth, we questioned whether or not the natural fungal comb environment provides the necessary metal ions for Fenton chemistry.^47^ To answer this question, we analyzed the element composition of fungus comb, gut fluids of termite workers and soil samples derived from within and outside termite colonies from different locations using atomic emission spectrometry (ICPAES).^48^ All tested samples contained Al, Fe, and Ti as some of the most abundant main elements, in addition to significant amounts of Mn. However, elements important for growth (C, H, P, K, Ca, Mg) were low in all soil samples, with a particularly strong depletion of phosphorus, but potassium was enriched compared to comb and gut samples (for details, see doi:10.5281/zenodo.4431413: Table S13, Figure S23-30, Table S23, S24). Sequential ion extraction of soil samples was performed to analyze the soluble metal ion content, and only low concentrations of most metal ions were detectable.^49,50^ Although these findings indicate that fungus comb and gut environment accommodate larger amounts of insoluble Fe/Al-oxide-containing clay minerals, the nano- and microscopic surface areas could provide the necessary catalytic centers for Fenton-like redox chemistry.^51^

### Enzyme activity tests catalyzing degradation of lignin model compounds

We then questioned if enzymatic degradation of lignin or lignin-type model substances by *Termitomyces* is measurable using colorimetric assays or MS-based analytical tools.^52^ For a first test, we supplemented culture medium of *Termitomyces* sp. T153 with the pigment-based model substance Azure B, previously used to measure redox-activity of LPs due to its stability towards oxidative activity of MnPs. Monitoring the decolourization of Azure B over time revealed that *Termitomyces* started to degrade Azure B seven days after inoculation; an effect which became more pronounced with increasing biomass and age of the fungus culture. To evaluate if the degrading activity of secreted enzymes and/or H_2_Q-mediated Fenton-based chemistry was responsible for the degradation of Azure B, we tested both effectors separately and in combination. While quantifying the enzymatic effects caused technical challenges due to intrinsic light absorption of enzyme concentrates, H_2_Q-mediated Fenton chemistry clearly induced the degradation of Azure B within five to ten minutes compared to the control (Fenton reagents without H_2_Qs) (for details, see doi:10.5281/zenodo.4431413: Figure S19,S20). ^53^ We then evaluated whether or not laccase activity was detectable within the secretome using a syringaldazine-based assay and compared the activity to the reactivity of a commercial laccase from *Trametes versicolor*,^**54**^ but only residual laccase activity was detectable in enriched enzyme extracts derived from different Termitomyces culture compared to the positive control and thus was unlikely accountable for the degradation of Azure B. Lastly, we evaluated if *Termitomyces* exhibits MnP enzymatic activity, which is marked by the oxidation of Mn^2+^ to Mn^3+^ and the release of the highly reactive oxidant as a carboxylic acid chelate using a previously reported leukoberbelin blue test.^55^ As shown in Figure 8, leukoberbelin-containing *Termitomyces* cultures and cell-free culture supernatant resulted in the formation of the blue leukoberbelin complex within minutes, which indicated the formation of Mn^+3/+4^ species. When *Termitomyces* was grown on PDA plates containing both, elevated Mn^2+^ concentrations (200 - 500 μM) and indicator dye, the formation of blue-colored leukoberbelin-Mn^3+/4+^ complexes was detectable within a few days and longer incubation times resulted in macroscopic-sized MnO_x_ precipitates forming around fungal hyphae within 10-17 days (Figure 4C). We further confirmed the gene expression encoding for the putative MnP by reverse transcription polymerase chain reaction (RT-PCR) (for details, see doi:10.5281/zenodo.4431413: Figure S17,S18).

**Figure 8.**
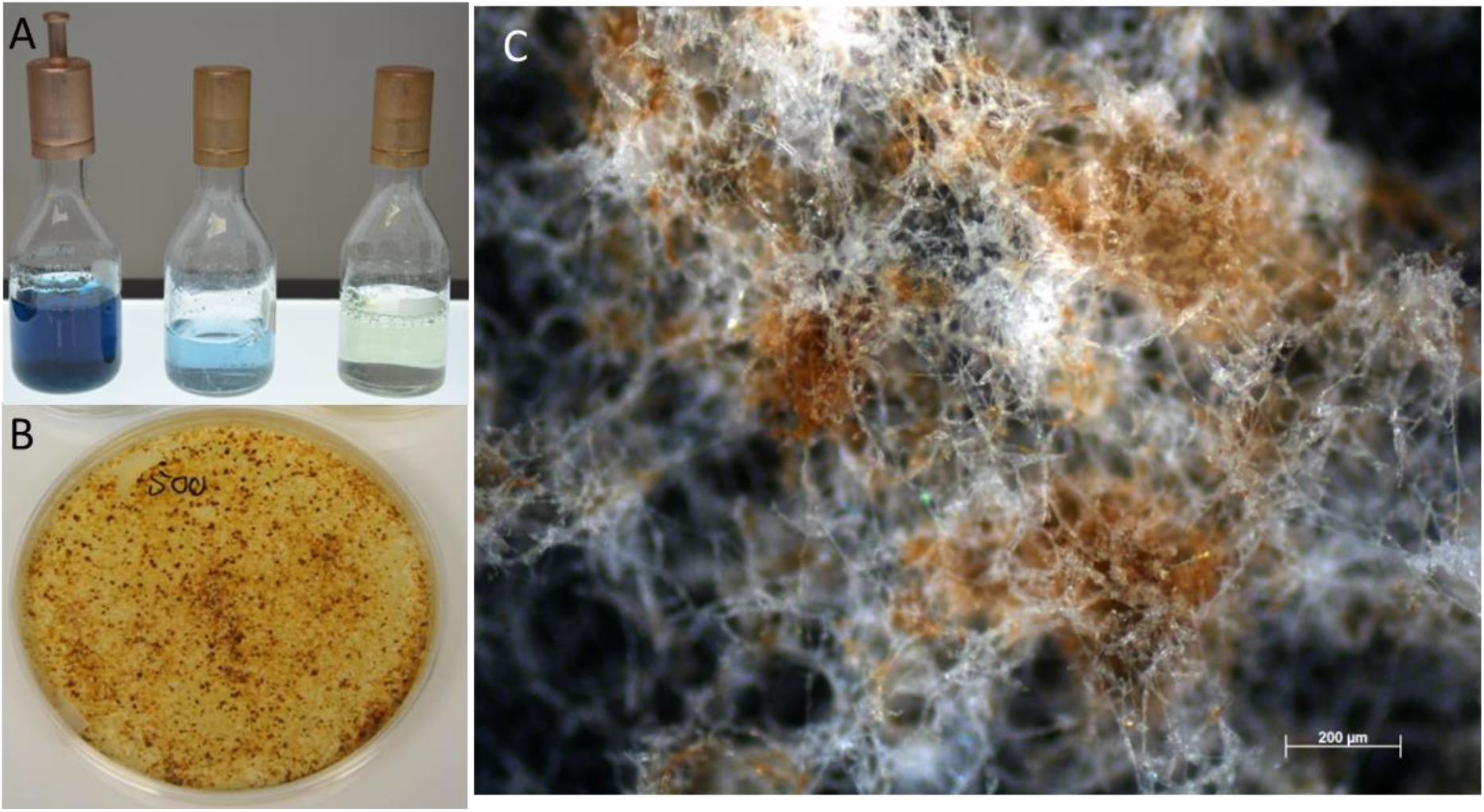
A) PDB containing Leukoberbelin blue (left to right: culture of *Termitomyces* sp. 153, cell-free supernatant, and PDB broth); B) *Termitomyces* sp. T153 cultivated on PDA containing 500 μM MnCl_2_ after 28 days and C) microscopic image of fungal mycelium after 24 d showing brown MnO_2_ deposits.

### Proteomic analysis

Building on our enzymatic studies and to link the observed activities with their putative enzymatic origin, we conducted a liquid-chromatography (LC)-MS/MS based proteomic analysis of secreted enzymes of *Termitomyces* culture supernatants, which were prepared in two different buffer systems (NaOAc, pH 4.5/KH_2_PO_4_, pH 6.5). Overall, a total of (255/303) secreted proteins were detectable, which were mostly assigned to fungal carbohydrate metabolism such as glucosidases, glucanases or chitinases (for details, see doi:10.5281/zenodo.4431413: Table S27-30). Interestingly, a potential lignin degrading aromatic peroxygenase (8^th^/13^th^) and one MnP (13^th^/11^th^) ranked amongst the top 15^th^ most abundant protein sequences, while two other yet unassigned peroxidases were also detectable (31^st^, 141^th^/17^th^, 142^nd^) albeit with lower abundance. In total five laccases were also detectable in minor abundances (starting from 76^th^/99^th^).

## Discussion

In the two major fungus-growing termite genera *Macrotermes* and *Odontotermes*, the decomposition of plant biomass by the fungal cultivar *Termitomyces* is based on the intricate interactions between the pre-digestive gut passage and the external fungus comb bioreactor. Although a series of studies have elaborated on the functional roles of *Termitomyces* in plant biomass degradation,^1,2,3^ experimental insights into the biochemical mechanisms necessary for plant biomass degradation have remained sparse.

### Which ligninolytic enzymes are produced by *Termitomyces*?

Our OMICs-based analysis clearly shows that *Termitomyces* has the capacity to produce a specific set of extracellular lignocellulose‐degrading enzymes, such as laccases (Lac), (aryl)-alcohol oxidases (AA3), and a manganese peroxidase (MnP, AA),^56^ all of which generate diffusible extracellular oxidants (superoxide O_2_^−^, hydroxyl radicals OH•, H_2_O_2_, redox-active Mn^3+/4+^ species or phenoxy-radicals) that oxidize the aromatic polymeric 3D structure of lignin (Figure 1D and 9). It is particularly intriguing that *Termitomyces* encodes on average for 16 different laccases that are differentially transcribed and might differ in their reactivity and substrate spectrum. And although laccases are considered not to be essential for lignin degradation, their presence likely assists in partial oxidation of phenolic and non-phenolic aromatic moieties that facilitate further fragmentation and depolymerization (Figure 9). Here, it is also worth highlighting that encoded and in culture secreted (aryl)-alcohol oxidases (AA3) are able to efficiently oxidize and cleave β-ether units present within lignin substructures via single electron transfer reactions. Our study also provides conclusive genomic and biochemical evidence that *Termitomyces* secretes a highly active manganese peroxidase MnP, an enzyme that oxidizes Mn^2+^ to the more reactive Mn^3+/4+^ and is known to be essential for extracellular degradation mechanisms. While none of these enzymes alone are capable of degrading lignin, their combined enzymatic action should allow for the partial depolymerization of lignin that is necessary for other enzymes to overcome the physical barrier of this complex phenolic polymer to initiate further degradation.

**Figure 9.**
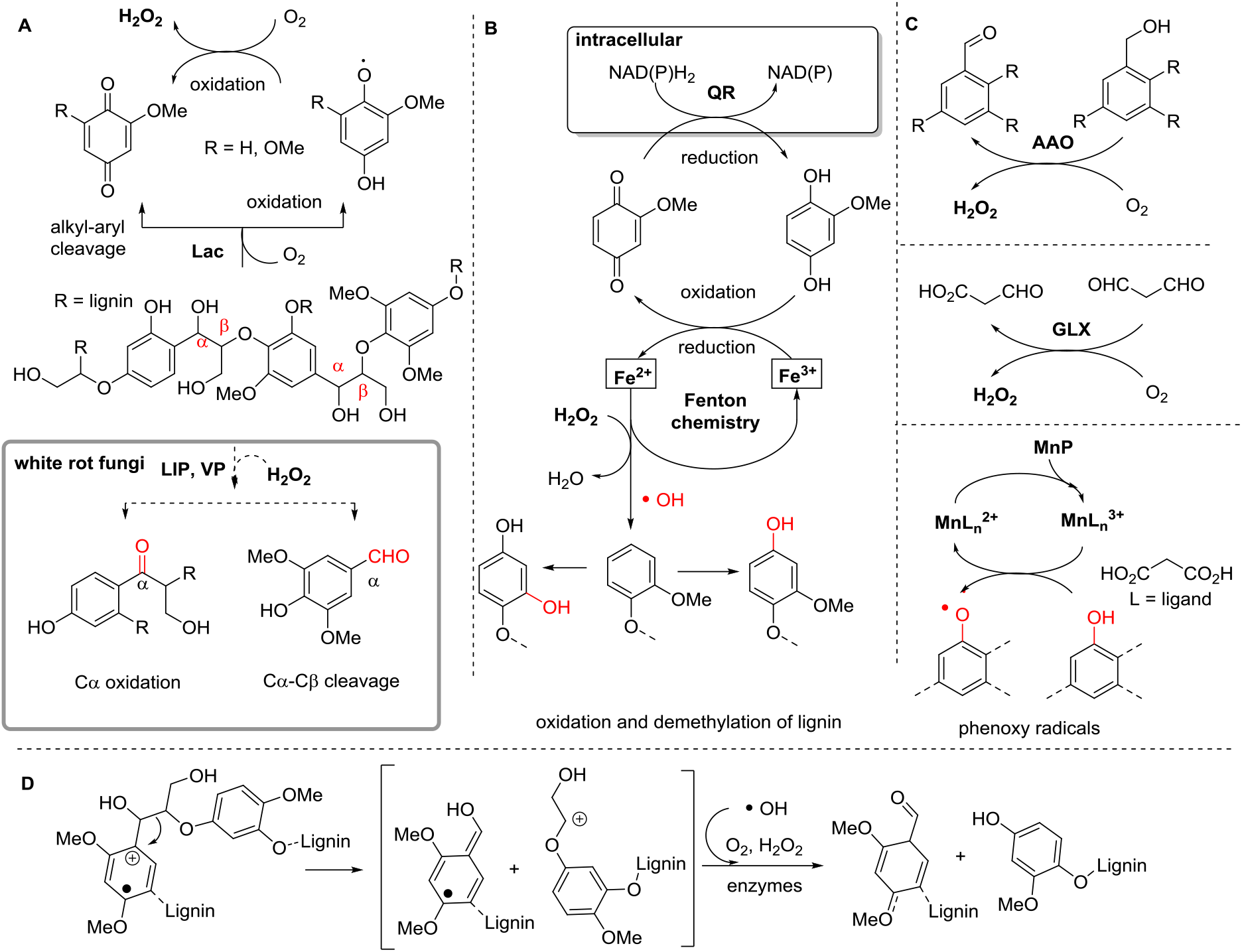
Lignin modifications and oxidation pathways by *Termitomyces*. A) Schematic depiction of lignin oxidation via Lac activity in *Termitomyces* and degradation by LIP and VP typically found in white-rot fungi (not found in *Termitomyces*, bold box). B) Oxidation and oxidative demethylation of lignin substructures by 2-MH_2_Q-catalyzed Fenton chemistry via the formation of short-lived hydroxyl radicals and regeneration of H_2_Q by (intracellular) benzoquinone reductases. C) Formation of H_2_O_2_ by (aryl)-alcohol oxidases (AAO) and glyoxal oxidases (GLX). D) Phenoxy radical formation catalyzed by the action of MnP ad oxidative C-C cleavage of lignin substructures by radicals and/or enzymatic processes.

### Does Fenton-chemistry play a role?

Following up on the idea that *Termitomyces* utilises Fenton-chemistry for biomass degradation, we evaluated the presence and absence of metabolic and enzymatic factors necessary to drive the radical process and identified collective evidence that *Termitomyces* employs Fenton-chemistry to degrade lignin-rich biomass by secretion of high levels of (extracellular) H_2_O_2_ and the production of H_2_Qs that reduce Fe^3+^ to Fe^2+^ necessary for Fenton chemistry. For the first time, we also document that the fungal metabolite 2-MH_2_Q (**19**) acts as a redox-shuttle for Fenton chemistry and induces the formation of Fe^2+^ similar to 4,5-DMH_2_Q and 2,5-DMH_2_Q.^20,34^ Although 2-MH_2_Q (**19**) lacks one −OCH_3_ compared to 2,5- or 2,6-DMH_2_Q, only a moderate decrease in activity was observed that likely correlates only with a minor shift in the reduction potential. Considering that Fenton chemistry produces several strong oxidants, we evaluated the influence of different dicarboxylic acids commonly secreted by fungal species, and monitored their influence on the H_2_Q-based reduction of Fe^3+^-complexes. Thus, it is reasonable to hypothesize that *Termitomyces* is capable of actively altering the redox properties of metal complexes within the surrounding fungal hyphae to protect itself against high oxidative stress. Genomic and transcriptomic evidence further suggests that *Termitomyces* produces two benzoquinone reductases that may reduce MQ to MH_2_Q and thereby close the H_2_Q/Q-based redox shuttle cycle.

Finally, our studies provide evidence that the natural environment contains sufficient amounts of iron and manganese to pursue Fenton chemistry,^51^ which are comparable to previous soil remediation studies.^47,49^ However, most metals are likely present as insoluble Fe/Al/Ti/Mn-oxides which let us to hypothesize that either microscopic Fe-rich minerals might serve as catalytic surface for the degradation of organic material,^51^ and/or the presence of chelating agents and reducing conditions might allow for the local formation Fe^2+/3+^.

## Conclusions

Collectively, our genomic, transcriptomic and proteomic studies document that *Termitomyces* harbours an enormous enzymatic repertoire to cope with the challenging task of depolymerizing the lignocellulose polymer to access cellulolytic components of the provided plant biomass, but lacks the genetic basis for the production of highly oxidizing versatile peroxidases that are known to be capable of oxidizing recalcitrant lignin parts. Furthermore, our chemical studies support the notion that the combined action of enzymatic degradation and Fenton chemistry are the key fungal contributions to the process of plant biomass decomposition, and Fenton reactions may, in part, complement the missing enzymatic capabilities. Whether or not symbiotic and lignocellulolytic bacteria^**57**^ present within the comb might also contribute and complement the fungal lignin-degradation capabilities is topic of current investigations and will further elaborate on the question why the *Termitomyces*-termite symbiosis has become the most successful path for the termite cultivar.

## Material and Methods

### Genome sequencing and processing

DNA was extracted from laboratory-grown heterocaryotic *Termitomyces* strains T112 and T153 and Genome sequences were produced at LGC Genomics (Berlin, Germany) using Illumina MiSeq V3 platform with 300bp paired-end reads and approx. 12 million read pairs per sequencing. All library groups were demultiplexed using the Illumina bcl2fastq 2.17.1.14 software (folder ‘RAW’, ‘Group’ subfolders). Up to two mismatches or Ns were allowed in the barcode read when the barcode distances between all libraries on the lane allowed for it. Sequencing adapters were clipped from all raw reads and reads with final length < 20 bases were discarded. Afterwards reads were quality trimmed by removal of reads containing more than one N, deleting reads with sequencing errors, trimming of reads at 3’-end to get a minimum average Phred quality score of 10 over a window of ten bases and discarding reads with final length of less than 20 bp. From the final set of reads, FastQC reports were created for all FASTQ files. Prior to annotation, the genomes were soft masked with RepeatMasker 4.0.9.^58^ RNAseq data was mapped to the genomes with STAR 2.7.3a ^59^ and used to train the Augustus gene predictor with Braker 2.1.5.^60^ Finally, the genomes T112 and T153 were annotated with Augustus 3.3.3.^61^ Protein and mRNA hints were used for the annotation (for details, see doi:10.5281/zenodo.4431413).

### RNA sequencing

RNA was obtained from mycelium of *Termitomyces* strains T153 and T112 cultivated on different growth media for 10 days at room temperature. Mycelium was harvested by scraping it from agar plates with a scalpel, freezing it in liquid nitrogen and storing it at −80 °C until RNA extraction. RNA extracts underwent 100bp paired-end BGISeq-500 sequencing with BGI (Hong Kong) (for details, see doi:10.5281/zenodo.4431413).

### RNAseq data acquisition and processing

RNAseq data for fresh comb (SRR5944783), old comb (SRR5944781) and nodules (SRR5944782) of *Termitomyces* strains from *Macrotermes* colony Mn156 were downloaded from the European Nucleotide Archive.^62^ The raw RNAseq data were mapped to the annotated genes of T153 using HiSat2 with spiced alignments disabled (Version 4.8.2).^63^ Transcript abundance was then estimated using HTSeq-count (Version 0.11.2).^64^ Count data from HTSeq were imported into R (R Core Team, 2018) using the “DESeq2” package (Version 1.22.2).^65^ Genes with low transcript abundance (<10) were filtered out and the remaining genes log10 transformed.^66^ A heatmap for the identified redox enzymes was generated using the “pheatmap” package (Version 1.0.12)^67^ in R (R Core Team, 2018)^68^ with color schemes generated by “viridis” (Version 0.5.1).^69^ (for details, see doi:10.5281/zenodo.4431413).

### CAZY Analysis

Identification of CAZymes in the predicted proteomes of *Termitomyces* and other Basidiomycetes strains was performed using a local installation of the dbCAN2 server^70^ and all three included tools (HMMER, DIAMOND, and Hotpep searches against the databases included in dbCAN2. For a reliable analysis, we kept only matches that were independently identified by at least two of three annotation strategies and only genes and transcripts classified by their substrate target and thus putative enzymatic functions. EC numbers were assigned using peptide-based functional annotation (www.cazy.org) (for details, see doi:10.5281/zenodo.4431413).^3,7^

### GC-MS analysis

The fungal isolates *Termitomyces* sp. P5 and T153 were cultivated on solid media containing different carbon sources. GC-MS analyses of biosamples were carried out with an Agilent (Santa Clara, USA) HP 7890B gas chromatograph fitted with a HP5-MS silica capillary column (30 m, 0.25 mm i. d., 0.50 μm film) connected to a HP 5977A inert mass detector (for details, see (for details, see doi:10.5281/zenodo.4431413).

### Activity studies on *Termitomyces* sp. T153

Detection and quantification of H_2_O_2_ in culture medium of *Termitomyces* sp. T153 was performed using a fluorimetric hydrogen peroxide assay kit (Sigma-Aldrich) (for details, see doi:10.5281/zenodo.4431413).

### Detection of hydroxyl radicals

Concentrations of hydroxyl radicals were measured using a fluorometric assay based on the reaction with terephthalic acid (TPA) yielding the fluorescent oxidation product hydroxy-terephthalic acid (hTPA) (for details, see doi:10.5281/zenodo.4431413).

### Ferrozin assay

Fe^2+^-concentrations were evaluated using a standardized Ferrozin-assay (for details, see doi:10.5281/zenodo.4431413).

### Proteomic Analysis

*Termitomyces* sp. T153 was cultured in Potato-Dextrose broth (25 mL) for 12 days (20 °C, 150 rpm) and secreted enzymes collected and digested according to standardized protocol (for details, see doi:10.5281/zenodo.4431413). LC-MS/MS analysis was performed on an Ultimate 3000 nano RSLC system connected to a QExactive Plus mass spectrometer (both Thermo Fisher Scientific, Waltham, MA, USA). Tandem mass spectra were searched against the UniProt database of *Termitomyces sp.* J132 (https://www.uniprot.org/proteomes/UP000053712;2019/11/04) using Proteome Discoverer (PD) 2.4 (Thermo) and the algorithms of Mascot 2.4 Sequest HT (version of PD2.2), and MS Amanda 2.0 (for details, see doi:10.5281/zenodo.4431413).

### Protein analysis and activity tests

Details regarding laccase and MnP activity tests are deposited under: doi:10.5281/zenodo.4431413).

## ASSOCIATED CONTENT

### Supporting Information

Supporting Information can be accessed free of charge at Zenodo: https://doi.org/10.5281/zenodo.4431413 and contain information regarding culture conditions, isolation procedures, structure elucidation, activity assays, expression level data, CAZY counts, and proteomic hit list.

## AUTHOR INFORMATION

### Author Contributions

The manuscript was written with contributions from all authors. All authors have approved the final version of the manuscript.

### Notes

### Conflict of Interest

There are no conflicts of interest to declare.

## ACKNOWLEDGMENT

Funded by the Deutsche Forschungsgemeinschaft (DFG, German Research Foundation) Project-ID 239748522 – SFB 1127 (project A6) to CB and CRC/TR 124 FungiNet, project number 210879364 (project A1 and Z2) to AB and OK. The Danish Council for Independent Research (DFF - 7014-00178) and a European Research Council Consolidator Grant (771349) to MP. Help with microscopy pictures by David Zopf is greatly acknowledged (SFB 1127/2 ChemBioSys – project number 239748522 (project Z). CG and NGC acknowledge the financial support from the state budget of the Slovenian Research Agency (Research Project J4-2549, Research Programs P1-0198 and P1-0170, Infrastructural Centre Mycosmo).

